# Chromosomal-level genome assembly of *Populus davidiana*

**DOI:** 10.1101/2023.07.11.548481

**Authors:** Liyang Chen, Yuanzhong Jiang, Tingting Shi, Changfu Jia, Zhiqin Long, Xinxin Zhang, Yupeng Sang, Jianquan Liu, Jing Wang

## Abstract

Despite the population structure and genetic differentiation of *P. davidiana* has been reported, little is known about the *P. davidiana* genome characterization with the genomes assembled at the chromosome level. As one of the most widely distributed and ecologically important tree species in China, *P. davidiana* is an excellent resource to understand population adaption to climate and environment change. Here we present a high-quality assembly based on long-read sequences and Hi-C data. This assembly is assembled into 19 contiguous chromosomes which provides a powerful tool for future association studies.

## Introduction

Increasingly dramatic climate change is emerging as a major threat to biodiversity, with rapid fluctuations in temperature and precipitation changing the suitability of specific areas. A large difference among environment and physiology can lead to changes in species ranges, population declines, and extinction. However, some species that are difficult to migrate have to survive only by adapting to rapidly changing environments. Species with sustainable variation in environment-related traits are most likely to have the ability to adapt to rapidly changing environments in order to counteract the pressures of drastic changes in the environment. Therefore, identifying potential adaptive alleles associated with the environment and demonstrating their distribution can accurately predict species responses to future environmental conditions and help mitigate climate change. Since multiple generations of genetic lineages are difficult to obtain for many wild non-model organisms, genomic data and environmental association analysis provide a different perspective for understanding adaptive evolutionary processes and assessing the future fitness of different populations. Here, we sequenced and *de novo* assembled a high-quality chromosome-level genome for *P. davidiana*, which is one of the most widely distributed and ecologically important tree species in China, aiming to provide a basic data resource for further genome-wide environmental association studies.

## Results and Discussion

### Genome Assembly

A preliminary genomic survey uncovered the estimated genome size of *P. davidiana* is ∼438 Mb and the heterozygosity rate is ∼1.6%. To obtain a high-quality genome assembly, we generated 47.37 Gb (∼94×) of Nanopore long-read sequences, 26.06 Gb (∼150×) of Illumina short-read data and 67.57 Gb (∼148×) of Hi-C reads. Contigs were initially assembled using Nanopore data and corrected by Illumina short reads. After removing the potential duplicate haplotypes as well as contaminated sequences in the genome, the resulting assembly consisted of 377 contigs, with a contig N50 of 2.35 Mb and a maximum length of 11.30 Mb (Table 1). These contigs were then scaffolded and assigned to their chromosomal positions using Hi-C data. The final assembly consists of 38 scaffolds, which spans 441.14 Mb in total, and with 98.23% anchoring to 19 pseudo-chromosomes with sizes ranging from 13.30 to 61.91 Mb (Table 2). The high completeness of this assembly was evidenced by a 98.11% Benchmarking Universal Single Copy Orthologs (BUSCO) recovery score (Table 3), indicating the continuity and quality of the assembled genome.

**Table 1.** Statistics of assembled contigs for *P. davidiana*.

| Type | Length | Number |
| --- | --- | --- |
| Total | 441,103,287 | 377 |
| Longest | 11,300,200 | - |
| N50 | 2,353,076 | 52 |
| N60 | 1,769,536 | 74 |
| N70 | 1,364,333 | 103 |
| N80 | 9,38,808 | 141 |
| N90 | 520,015 | 202 |

**Table 2.** Scaffolding of contigs based on Hi-C data.

| Peudo-chromosome | Length (Contig number) |
| --- | --- |
| LG01 | 61,911,085 (56) |
| LG02 | 28,485,360 (29) |
| LG03 | 27,480,174 (16) |
| LG04 | 26,170,615 (33) |
| LG05 | 25,594,639 (17) |
| LG06 | 25,427,964 (24) |
| LG07 | 23,487,940 (15) |
| LG08 | 22,866,961 (29) |
| LG09 | 22,154,157 (17) |
| LG10 | 22,082,649 (25) |
| LG11 | 21,848,605 (14) |
| LG12 | 17,912,890 (6) |
| LG13 | 17,603,465 (15) |
| LG14 | 16,709,382 (11) |
| LG15 | 16,025,600 (15) |
| LG16 | 15,153,166 (7) |
| LG17 | 14,703,232 (14) |
| LG18 | 14,362,361 (9) |
| LG19 | 13,299,919 (9) |
| Total | 433,280,164 (361) |

**Table 3.** Statistics of BUSCO evaluation for genome assembly.

| BUSCO | Number | Percentage |
| --- | --- | --- |
| Complete BUSCOs (C) | 1,349 | 98.11% |
| Complete and single-copy BUSCOs (S) | 1,032 | 75.05% |
| Complete and duplicated BUSCOs (D) | 317 | 23.05% |
| Fragmented BUSCOs (F) | 6 | 0.44% |
| Missing BUSCOs (M) | 20 | 1.45% |
| Total BUSCO groups searched | 1,375 | 100.00% |

### Genome annotation

Our assembly indicated that 176.9 Mb (40.10%) of the assembled *P. davidiana* genome consisted of repeated regions (Table 4). Long terminal repeats (LTRs) were the most abundant annotations, making up 22.58% of the genome, with *Gypsy* and *Copia* elements accounting for 12.59% and 5.07% of the genome, respectively.

**Table 4.** Statistics of repeat sequences in *P. davidiana* genome.

| TE type | Total size (bp) | Percentage of genome (%) |
| --- | --- | --- |
| **Class I: Retrotransposon** | 101,053,451 | 22.91 |
| **LTR Retrotransposon** | 99,623,383 | 22.59 |
| *Copia* | 22,362,310 | 5.07 |
| *Gypsy* | 55,524,652 | 12.59 |
| unkonwn | 21,736,421 | 4.93 |
| **LINE** | 1,430,068 | 0.32 |
| **Class II: DNA transposon** | 51,912,086 | 11.77 |
| Helitron | 24,603,053 | 5.58 |
| CACTA | 6,586,911 | 1.49 |
| Mutator | 14,365,370 | 3.26 |
| PIF_Harbinger | 1,408,202 | 0.32 |
| Tcl_Mariner | 1,362,230 | 0.31 |
| hAT | 3,586,320 | 0.81 |
| **Unclassified** | 23,913,435 | 5.42 |
| **Total** | 176,878,972 | 42.10 |

We masked repeated regions and proceeded to annotate the genome using a comprehensive strategy combining homology-based, transcriptome evidence-based and *ab initio* gene prediction. In total, 38,244 gene models were identified, with an average CDS length of 1.21 kb and an average of 4.93 exons per transcript (Table 5). We used BUSCO to evaluate the quality of our gene annotation and found that 1,588 out of the 1,614 (98.39%) highly conserved core proteins in the Embryophyta lineage were present in our gene annotation, of which 1,350 (83.64%) were single-copy genes and 238 (14.75%) were duplicated. For the remaining conserved genes, 5 (0.31%) had fragmented matches and 21 (1.30%) were missing.

**Table 5.** Statistics of protein-coding genes.

| Feature | number/length |
| --- | --- |
| Gene number | 38,244 |
| Average gene length (bp) | 3950.91 |
| Mean exons number per mRNA | 4.93 |
| Mean CDS number per mRNA | 4.79 |
| Average CDS length (bp) | 1126.26 |
| Average intron length (bp) | 2161.84 |

Functional annotation confirmed that 95.71% of these genes could be assigned to at least one of the databases TrEMBL, Swiss-Prot, NR, Pfam, Interproscan, GO or KEGG (Table 6).

**Table 6.** Statistics of protein-coding gene functional annotation.

| Database | Number | Percentage (%) |
| --- | --- | --- |
| Pfam | 26,651 | 69.69 |
| Interproscan | 34,569 | 90.39 |
| KEGG | 11,146 | 29.14 |
| NR | 35,405 | 92.58 |
| Swiss-Prot | 27,752 | 72.57 |
| KOG | 31,686 | 82.85 |
| COG | 12,724 | 33.27 |
| Tremble | 36,006 | 94.15 |
| GO | 28,276 | 73.94 |
| Unannotated | 1,641 | 4.29 |

## Methods

### Plant materials and genome sequencing

Fresh young leaves of *P. davidiana* were harvested from wild in Hanzhong, Shanxi, China. Genomic DNA was extracted with a standard CTAB method. For the Illumina short-read sequencing, paired-end libraries with insert sizes of 350 bp were constructed and sequenced using an Illumina HiSeq X Ten platform. For the long-read sequencing, the genomic libraries with 20-kbp insertions were prepared and sequenced using the PromethION platform of Oxford Nanopore Technologies (ONT). For the Hi-C experiment, approximately 3 g of fresh young leaves of the same *P. davidiana* accession was ground to powder in liquid nitrogen. Hi-C libraries were then constructed following the standard protocol and sequenced on an Illumina NovaSeq platform. To assist gene annotation, total RNAs from fresh young leaves, roots and stems were extracted with CTAB (Rogers and Bendich 1985) procedure to prepare RNA-seq libraries, which were subsequently sequenced on Illumina HiSeq X Ten platform.

### Genome assembly and pseudo-chromosome construction

The Illumina short reads were first used to estimate the genome size of *P. davidiana* via a 17-bp k-mer frequency analysis with Jellyfish (v2.3.0)^1^. NextDenovo (v2.0-beta.1, https://github.com/Nextomics/NextDenovo) was then performed to generate the preliminary sequence assembly based on the Nanopore long reads. The quality-controlled long reads were first corrected via a self-align method using NextCorrect with parameters “reads_cutoff=1k, seed_cutoff=30k”, and then assembled using NextGraph with default parameters. To improve the accuracy of the draft assembly, two-step polishing strategies were applied. The first step included three rounds of polishing by Racon v1.3.1^2^ based on the corrected ONT long reads. The second step included four rounds of polishing by Nextpolish v1.0.5^3^ based on cleaned Illumina short reads. The redundant sequences were subsequently removed using the perge_haplotigs v1.1.1^4^ software and the final contig-level assembly were checked for DNA contamination by searching against the NCBI non-redundant nucleotide database (Nt) using BLASTN, with an E-value cutoff of 1e-5. Finally, the completeness of the genome assembly was assessed by BUSCO (v4.0.5, embryophyta_odb10 download at 16-Oct-2020)^5^ with default settings.

For chromosome-level scaffolding, the Hi-C reads were first filtered by fastp v0.20.0^6^. The clean reads were then aligned onto the contig-level assembly by bowtie2 v2.3.2^7^ with parameters ‘-end-to-end, -very-sensitive -L 30’. The quality of Hi-C data was evaluated by HiC-Pro v2.11.4^8^ and only valid interaction pairs were retained for further analysis. Scaffolds were clustered, ordered and oriented onto 19 pseudo-chromosomes using LACHESIS^9^ with parameters CLUSTER MIN RE SITES=100, CLUSTER MAX LINK DENSITY=2.5, CLUSTER NONINFORMATIVE RATIO=1.4, ORDER MIN N RES IN TRUNK=60, ORDER MIN N RES IN SHREDS=60. The placement and orientation errors that exhibit obvious discrete chromosome interaction patterns were then manually adjusted

### Repeat and gene annotation

For transposable element annotation, the pipeline Extensive *de-novo* TE Annotator (EDTA v1.9.3^10^) was used to identify known repeats in the *P. davidiana* genome. Prior to gene prediction, RepeatMasker v4.1.0^11^ was performed to mask the whole genome sequences based on the EDTA-constructed TE library.

Three distinct strategies comprising *ab initio* prediction, homology search and expression evidence were combined to generate the predicted gene models in the *P. davidiana* genome. For transcriptome-based prediction, RNA from three tissues (leaf, root and stem) was isolated and the clean RNA-seq data, processed by Trinity v2.8.4^12^ with default parameters were used for gene annotation. For homology-based gene prediction, protein sequences from six sequenced plants (*Populus trichocarpa, Populus euphratica, Salix brachista, Salix purpurea, Arabidopsis thaliana* and *Vitis vinifera*) were initially aligned to the *P. davidiana* genome using TBLASTN (ncbi-BLAST v2.2.28). The homologous genome sequences were then aligned against the matching proteins using Genewise v2.4.1^13^. For *ab initio* gene prediction, AUGUSTUS v3.3.2^14^ was employed with default parameters after incorporating the transcriptome-based and homology-based evidence for gene model training. Finally, we combined the gene annotation results from all homology-based, *ab initio* and transcriptome-based predictions using EvidenceModeler v1.1.1^15^ to produce a consensus protein-coding gene set.

Functional annotation of the predicted gene models was based on comparison with the NCBI nonredundant protein database (NR), Swiss-Prot, TrEMBL, COG and KO databases with a minimal e-value of 1e-5. Protein domains and functions as well as gene ontology (GO) terms were assigned to the annotated genes using InterProScan (v5.32-71.0)^16^.

